# On the edge of criticality: strength-dependent perturbation unveils delicate balance between fluctuation and oscillation in brain dynamics

**DOI:** 10.1101/2021.09.23.461520

**Authors:** Yonatan Sanz Perl, Anira Escrichs, Enzo Tagliazucchi, Morten L. Kringelbach, Gustavo Deco

## Abstract

Despite decades of research, there is still a lack of understanding of the role and generating mechanisms of the ubiquitous fluctuations and oscillations found in recordings of brain dynamics. Here, we used a strength-dependent perturbative framework to provide a causal mechanistic description of how human brain function is perched at the delicate balance between fluctuation and oscillation. Applying local strength-dependent perturbations and subsequently measuring the perturbative complexity index clearly demonstrates that the overall balance of brain dynamics is shifted towards fluctuations for providing much needed flexibility. Importantly, stimulation in the fluctuation regime modulates specific resting state network, thus providing a mechanistic explanation of experimentally reported brain dynamics. Furthermore, this framework generates specific, testable empirical predictions for human stimulation studies using strength-dependent rather than constant perturbation. Overall, the strength-dependent perturbative framework demonstrates how the human brain is poised on the edge of criticality, between fluctuations to oscillations, allowing for maximal flexibility.

## Introduction

Already at the birth of neuroscience, a deep problem emerged: namely that local and global recordings from inside and outside the brain show very complex fluctuating and oscillating patterns of brain activity (Biswal et al., 1995; Destexhe & Contreras, 2006; M. Fox, 2005; Mantini et al., 2007; M. Raichle & Mintun, 2006). This gave rise to the fundamental question of the importance of synchronous or asynchronous local dynamics as the origin of the dynamical behaviour of brain states (Goldman et al., 2019, 2020). This distinction is important since fluctuating versus oscillating scenarios emerge from very different principles, such as when pressure fluctuations in the water surface give rise to long range structures like waves (Phillips, 1957), which is different from when entrainment synchronises oscillating biological clocks with the availability of light (Mondragón-Palomino et al., 2011). In global brain dynamics, a purely fluctuating scenario will give rises to patterns formed due to noise correlations, whereas a purely oscillatory regime would produce patterns arising mainly from cluster synchronisation. In both cases, the activity is shaped by the underlying brain anatomy but the generating principles are clearly different. Even more, the asynchronous, irregular background dynamics have been associated with conscious, responsive brain state, while synchronisation and regular dynamics have been linked with reduced states of conscious awareness (Goldman et al., 2019).

Deco & Kringelbach have proposed a novel framework based on recent results showing turbulence in the brain dynamics, which is based on quantifying the level of local synchronisation in whole-brain activity (Deco, Kemp, et al., 2021; Deco, Perl, et al., 2021; Deco & Kringelbach, 2020). The brain as a non-fluid system has been shown to be turbulent in the sense defined by the coupled oscillator framework of Kuramoto (Kawamura et al., 2007; Kuramoto, 1984). Furthermore, in terms of brain function, a key desirable property of turbulence is its mixing capability due to the energy/information cascade, which has been shown to be highly efficient across scales as shown by the power law discovered by Kolmogorov (Kolmogorov, 1941b, 1941a). Note that for these power laws, the flow is not provided by billions of molecules in a fluid but by the flow of information coming from the interplay and mutual entrainment in billions of neurons underlying brain dynamics.

Further to these empirical observations, the relevance of whole-brain computational model within this analytical framework was demonstrated based on the fact that brain dynamics can be accurately modelled by a system of coupled non-linear Stuart-Landau working in a turbulent regime (Deco, Perl, et al., 2021; Deco & Kringelbach, 2020), which integrates anatomy and local dynamics (Deco et al., 2018; Deco, Kringelbach, et al., 2017; Jobst et al., 2017; Saenger et al., 2017). As it happens, this whole-brain model is suited to resolve the question in hand, since it naturally describes the transitions between noise, fluctuation and oscillation. The whole-brain system can produce three radically different regimes, simply by varying the local bifurcation parameter: 1) *Noise* regime – when the parameter is much less than zero, 2) fluctuating *subcritical* regime – when the parameter is just below zero; and 3) oscillatory *supercritical* regime – when the parameter is larger than zero. Previous research has shown that the three different scenarios of noise (Ghosh et al., 2008; Messé et al., 2014), subcritical (Deco, Kringelbach, et al., 2017; Deco & Jirsa, 2012; Ghosh et al., 2008; Hansen et al., 2015) and supercritical (Cabral et al., 2014) are equally able to fit the empirical neuroimaging data in terms of functional connectivity. However, it is not clear when fitting the whole-brain model to the more sensitive measure of turbulence which regimes are able to fit the empirical data.

In a complementary direction, empirical perturbations have proved to be an excellent approach to provide insights into the complexity of brain dynamics, such as the one proposed by Massimini and colleagues. The authors used transcranial magnetic stimulation (TMS) with electroencephalography (EEG) to demonstrate perturbation-elicited changes in global brain activity in the perturbative complexity index (PCI) between different brain states (wakefulness, sleep, anaesthesia and coma) (Casali et al., 2013; Ferrarelli et al., 2010; Massimini et al., 2005). The results showed, for example, that non-REM sleep is accompanied by a breakdown in cortical effective connectivity, where the stimuli rapidly extinguish and do not propagate beyond the stimulation site (Casali et al., 2013; Ferrarelli et al., 2010; Massimini et al., 2005). Based on the same strategy, previous experimental research demonstrated that stimulation with TMS in specific brain regions can differentially modulates specific networks (Santarnecchi et al., 2018; Tik et al., 2017). Computational approaches also demonstrated that *constant* perturbation can also provide clear insights into the complexity of brain dynamics (Deco et al., 2018; Goldman et al., 2020; Kunze et al., 2016).

To address the question of distinguishing between fluctuation and oscillations in shaping global brain dynamics, their origin and balance, we constructed and perturbed whole-brain models taking advantage of the recently proposed turbulence framework. Specifically, to disentangle the generative roles of the fluctuation (subcritical) and oscillations (supercritical) models, we created a global and local strength-dependent perturbational framework. This allowed us to observe the evolution of three perturbative sensitivity measures, susceptibility, information capacity (defined in the turbulence framework (Deco, Perl, et al., 2021)) and the well-known PCI, as a function of the applied global sustained perturbation. The stimulation in the fluctuation regime also allows us to provide a mechanistic explanation of experimentally reported brain dynamics showing the modulation of specific resting state networks when are perturbed. With this approach, we were able to show that using strength-dependent perturbations - instead of the classical flat, constant perturbations - provides the means for disentangling alternative model-based hypotheses about the underlying empirical dynamics. Our results show both how and why the human brain is poised on the edge of criticality, between fluctuations and oscillations, allowing maximal flexibility in information processing.

## Results

In this study, we were interested in revealing the underlying mechanisms of the different dynamical regimes available in the resting state (M. Fox, 2005; M. E. Raichle et al., 2001). We extended the well stablish perturbative approach to use strength-dependent, non-constant perturbation in a whole-brain model fitting the empirical data to provide a causal mechanistic explanation for disentangling fluctuating from oscillating regimes in the underlying empirical dynamics. **Figure 1** shows the details of our framework, which has two key ingredients: 1) a *model-based* approach which is probed with 2) varying levels of *strength-dependent perturbations*. The whole-brain model is based on the recent breakthrough of demonstrating turbulence (**Figure 1A**) in empirical neuroimaging data (**Figure 1B**). Turbulence is a property found in high-dimensional non-linear systems, where its mixing capability is crucial for giving rise to the efficient energy/information cascade, whereby large whirls turns into smaller whirls and eventually energy dissipation. Using this novel turbulence framework, we were able to determine the vorticity, i.e. the local level of synchronisation, allowing us to extend the standard global time-based measure of metastability to become a local-based measure of both space and time for capturing the essential features of the underlying brain dynamics.

**Figure 1.**
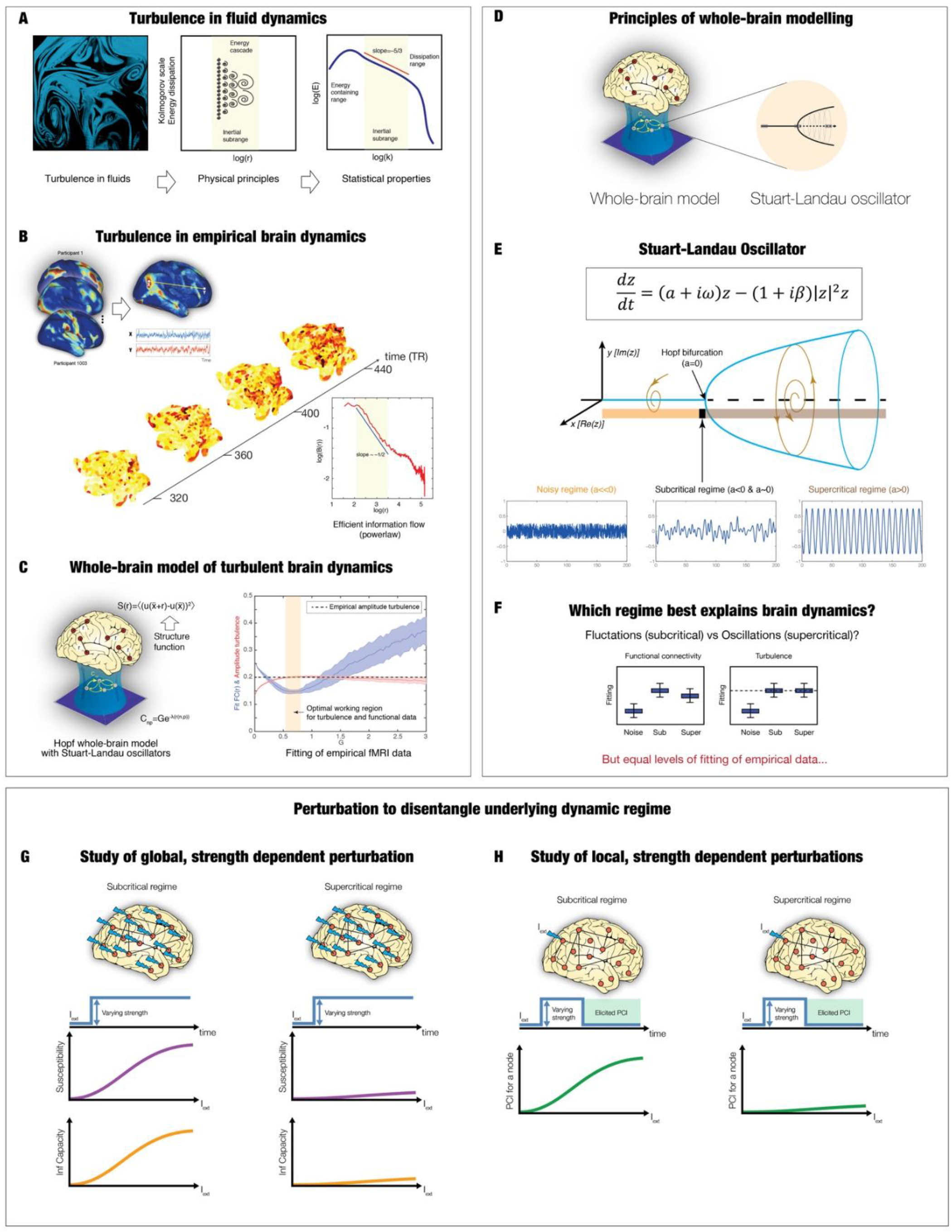
Overview of the framework. **A)** Turbulence provides a good description of the seemingly chaotic dynamics of fluids as first described by Leonardo da Vinci (Deco, Kemp, et al., 2021) (left panel drawing of turbulent whirls). The physical principles giving rise to turbulence are given by high-dimensional spacetime non-linear coupled systems. In turbulence, a fundamental property is its mixing capability which yields the energy cascade through turning large whirls into smaller whirls and eventually energy dissipation (middle panel). Furthermore, the turbulent energy cascade has been shown to be highly efficient across scales, as evidenced by a power law (right panel). **B)** Empirical brain dynamics was recently shown to exhibit turbulence (Deco & Kringelbach, 2020). The fMRI resting state analysis over 1000 healthy participants (left panel) shows the presence of highly variable, local synchronisation vortices across time and space (middle panel). Equally, the turbulent brain regime also gives rise to an efficient information cascade obeying a power law (right panel). **C)** Furthermore, Hopf whole-brain models (Deco, Kringelbach, et al., 2017) (left panel) were able to fit both turbulence and the empirical data at the same working point (right panel). **D)** We model brain activity as a system of non-linear Stuart Landau oscillators, coupled by known anatomical connectivity. **E)** The Stuart Landau equation (top panel) is suited for describing the transitions between noise and oscillation. By varying the local bifurcation parameter, a, the equation will produce three radically different regimes: Noise (a<<0), fluctuating subcritical regime (a<0 & a∼0) and oscillatory supercritical regime (a>0) (bottom panel). **F)** We evaluated the fitting capacity of the three model regimes in terms of functional connectivity and turbulence (with the dashed line showing the empirical level of turbulence). **G)** However, it is well-known that physical systems can be more deeply probed by perturbing them. Therefore, we used strength-dependent perturbations to disentangle the generative roles of the fluctuation (subcritical) and oscillations (supercritical) models. We observed the evolution of two key perturbative measures, susceptibility and information capacity, as a function of the applied global sustained perturbation. **H)** Finally, in order to generate experimentally testable hypotheses, we used local strength-dependent, non-sustained perturbations and measured the elicited dynamics in terms of the empirical perturbative complexity index (Casali et al., 2013). Specifically, we simulated 600 volumes with the perturbation active, and we then evaluated the evolution of the signals in the following 200 volumes without perturbation and computed the difference between the PCI after and before perturbation.

The Hopf whole-brain model can fit the complex spatiotemporal brain dynamics in terms of both functional connectivity and turbulence (**Figure 1C**). More generally, the Hopf whole-brain model integrates anatomical connections (Hagmann et al., 2007, 2010; Johansen-Berg & Rushworth, 2009) with local dynamics to explain and fit the emergence of global dynamics in empirical data (Deco, Cabral, et al., 2017; Deco, Kringelbach, et al., 2017; Freyer et al., 2012; Ipiña et al., 2020; Piccinini et al., 2021) (**Figure 1D**). For decades, brain signals have been recorded with a plethora of different techniques showing them to be combinations of at least three different regimes: noise, fluctuating, and oscillatory. The non-linear Stuart Landau oscillator is perfect for generating and testing these three regimes, given that the local bifurcation parameter in the equation governs the dynamics of each local brain region (**Figure 1E**). Indeed, by varying this parameter the Stuart-Landau equation will produce three radically different signals: 1) a noise signal resulting from Gaussian noise added to a fixed point when the parameter is much less than zero; 2) a fluctuating stochastically structured signal when the parameter is just below zero; 3) an oscillatory signal when the parameter is larger than zero. Technically, these three regimes are termed noise, subcritical and supercritical, respectively.

In the following, we show that the subcritical fluctuating and supercritical oscillatory regimes are equally able to fit the empirical data in terms of functional connectivity and turbulence (**Figure 1F**). Crucially, however, our framework includes the second ingredient of strength-dependent perturbation, which, as shown below, has allowed us to distinguish between the two regimes. We probe the model in two ways using both global (**Figure 1G**) and local strength-dependent perturbations (**Figure 1H**) and measuring the sensitivity of the system through quantifying the elicited susceptibility, information capacity and perturbative complexity index.

### Hopf whole-brain model of large-scale empirical neuroimaging data

We first investigated the ability of the three regimes to fit empirical data. We fitted whole-brain models of Stuart Landau oscillators in the three regimes to the large-scale neuroimaging resting state fMRI data from 1003 healthy human participants in the Human Connectome Project (HCP) (Van Essen et al., 2013). We extracted the timeseries in the Schaefer1000 parcellation (Schaefer et al., 2018), a fine-grained atlas that allowed quantified turbulence in empirical data (Deco & Kringelbach, 2020).

Previous Hopf whole-brain models have successfully fitted functional neuroimaging data with different acquisition parameters from many different neuroimaging setups (Donnelly-Kehoe et al., 2019; Ipiña et al., 2020; Perl et al., 2021) using a fluctuating regime with a local bifurcation parameter close to the bifurcation point. Here we aim to fit both functional connectivity and turbulence of functional neuroimaging data. In order to fit turbulence with the Stuart-Landau oscillator in the oscillatory supercritical regime, Kuramoto and colleagues (Kawamura et al., 2007) have shown that an extra parameter, the so-called shear parameter, is fundamental. Therefore, we extend the Hopf whole-brain model to use the appropriate formulation of the Stuart-Landau equation (see Methods) to be able to fit the data with the supercritical regime.

We explored the parameter space of varying the global coupling (G) and the shear parameter (β). The coupling parameter (G) scales the local fibre densities of the anatomical structural connectivity (see Methods) to capture the effectivity of the coupling by assuming a single global conductivity parameter. The shear parameter (β) acts similar to viscosity in fluid dynamics (Kuramoto, 1984) in that it is able to affect both the frequency and amplitude of the generated oscillations (Kawamura et al., 2007). Importantly, for the structural connectivity in the whole-brain model, we used a combination of exponential distance rule, EDR (Ercsey-Ravasz et al., 2013) and long-range connections, which improve and provide an excellent fit to the available dMRI tractography from humans (Deco, Perl, et al., 2021)(see Methods).

In order to fit the whole-brain model, we used the following observables: 1) the empirical mean level of amplitude turbulence, as the standard deviation of the Kuramoto Local order parameter (D), in fine parcellation, and metastability, as the standard deviation of the Kuramoto Global order parameter (M), in coarse parcellation; and 2) the grand average functional connectivity (FC) from the neuroimaging empirical data (see Methods). For measuring the level of fitting for each: 1) for turbulence/metastability measure, we computed the error (eD= abs(D_sim_-D_emp_)/(eM= abs(M_sim_-M_emp_)), i.e. by the absolute difference between the simulated and empirical amplitude turbulence and 2) for the functional connectivity, we computed Euclidean distance (eFC) between the simulated and empirical FC.

### Modelling results for fine-scale parcellation with 1000 regions

**Figure 2** shows the results of fitting the Hopf whole-brain model in the three different regimes (noise, fluctuating and oscillatory, see upper row) for the Schaefer1000 parcellation in terms of functional connectivity and turbulence. For each of these regimes, we defined a grid of the parameter space (G,β), where G is the coupling strength factor, i.e. the global scaling factor of regional connectivity and β, the shear parameter (see above and Methods). For each pair in the grid, the whole-brain dynamics were simulated 100 times, and we computed the level of fitting between amplitude turbulence (second row) and simulated and empirical FC (third row).

**Figure 2.**
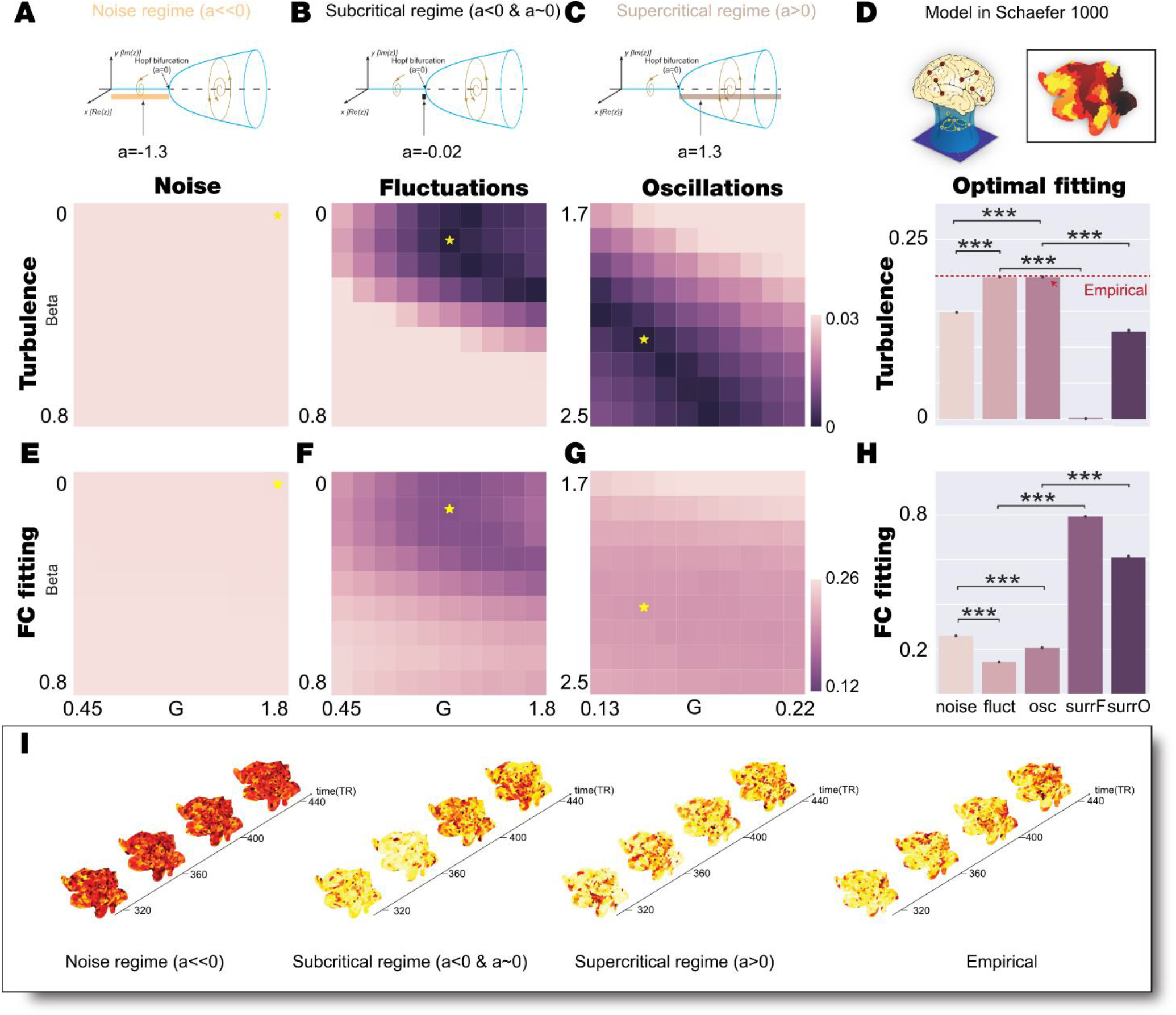
Model Schaefer1000 fitting of noise, fluctuations, and oscillatory for 1) Turbulence and 2) FC. **A-C)** We explored the bi-dimensional parameter space defined by β and G for noise, fluctuating and oscillatory regime (bifurcation parameter a=-1.3, a=-0.02 and a=1.3, respectively, indicated in upper row). We computed the level of amplitude turbulence error as the absolute difference between the empirical and simulated turbulence. Yellow stars indicate the (β, G) combination that reaches the lowest turbulence error in each regime. **D)** The upper subpanel shows the model fitting scheme in fine Schaefer1000 parcellation. The bottom subpanel displays the barplot that indicates the statistical distribution of the level of amplitude turbulence obtained by simulating 20 trials with 100 subjects for each model regime with the parameters set at the corresponding working point. We also display the results of two model-based surrogates created by increasing the shear parameter of each model regime. The red dashed line indicates the empirical level of amplitude turbulence averaged across participants. The subcritical, supercritical and empirical level of turbulence are not statistically different (Wilcoxon test, ns), the rest of the comparison are statistically significant (Wilcoxon test, P<0.001). **E-G)** We explored the bi-dimensional parameter space defined by β and G for noise, fluctuating and oscillatory regime computed the FC fitting as Euclidean distance between the empirical and simulated FC. Yellow stars indicate the (β, G) combination that reaches the lower turbulence error in each regime (the optimal working point obtained in panels A-C). **H)** The barplot indicates the statistical distribution of the FC fitting obtained by simulating 20 trials with 100 subjects for each model regime at the corresponding working point defined as the minimum turbulence error. We also display the results for the model-based surrogates. All comparisons are statistically significant (Wilcoxon, P<0.001). **I)** Visualization of the change of the local Kuramoto order parameter, R, in space and time reflecting amplitude turbulence in a single simulation at the optimal working point of each regime (noise, fluctuating and oscillatory cases) and one participant (empirical). This can be appreciated from continuous snapshots for segments separated in time rendered on a flatmap of the hemisphere.

We found the optimal fitting for turbulence for each of the three regimes, indicated with a star in the second row of **Figures 2A-C. Figure 2A** shows the best fit for the noise regime (a=–1.3) with optimal (G,β)=(1.8,0) as the absolute difference between the simulated and empirical turbulence D (here eD=0.0473). **Figure 2B** shows the best fit for the fluctuating regime (a= –0.02) with optimal (G,β)=(1.2, 0.1), which produces an excellent fit with eD= 4×10^−4^. **Figure 2C** shows the best fit for the oscillatory regime (a=1.3) with optimal (G,β)=(0.15, 2.2), which also produces an excellent fit with eD=3×10^−4^. We also computed the grid fitting for the FC for all three regimes (**Figures 2E-G**), with a star in the grid indicating the optimal fit of turbulence which is the criterium for selecting the optimal working point since this is a more sensitive measure. Note that these points do not correspond to the optimal fitting with FC in the three regimes.

In summary, both the fluctuating and oscillatory regimes are excellent for fitting the turbulence in the empirical data, while the noise regime is not. This is quantified in **Figure 2D**, which shows the statistical comparisons between optimal fitting (indicated with the stars in **Figures 2A-C**) of the three regimes with turbulence (repeated 20 times 100 simulations) and a horizontal line of D=0.1976 indicates the level of empirical turbulence.

The results show that the best working point for the noise regime is only giving mean D=0.1484, which is significantly worse than both fluctuating and oscillatory regimes (Wilcoxon p<0.001, compare noise and fluctuation; Wilcoxon p<0.001, compare noise with oscillatory in **Figure 2D**). On the other hand, the fitting of the data by the fluctuating and oscillatory regimes is excellent (mean D=0.1972 and mean D=0.1973 respectively) but not significantly different (Wilcoxon n.s, comparing second with the third bar in **Figure 2D**).

Further bolstering these findings, we also generated model-surrogates (see Methods) to compare with the corresponding optimal working point by setting the parameters of the model in the optimal working point but increasing β, which is known to suppress turbulence (Deco & Kringelbach, 2020). Hence, we produced two surrogate models: *surr_fluct* for the fluctuating model surrogates using *a=-0*.*02* and (G,β) =(1.2, 6) and *surr_osc* for the oscillatory model surrogates using *a=1*.*3*. and (G,β) =(0.15, 6). The results clearly show significant differences comparing with the level of turbulence fitting obtained by the optimal working point of the model in different regimes (Wilcoxon p<0.001, comparing fluctuation with *surr_fluct* and comparing oscillations with *surr_osc*).

Similarly, we fitted the whole-brain model with the functional connectivity by means of the Euclidean distance with the empirical. In **Figures 2E-G**, we show the fitting for (G,β) for the noise, fluctuating and oscillating regimes. We quantify the fit (using the optimal points from the turbulence fitting indicated with the stars in **Figures 2A-C**) in **Figure 2H**, which shows the statistical comparisons of the three regimes with functional connectivity (see Methods).

The results show that the best working point for the turbulence fitting for the noise regime is only giving a functional connectivity fitting mean ErrFC=0.2594, which is significantly worse than both fluctuating and oscillatory regimes (Wilcoxon p<0.001, comparing noise with fluctuations [mean ErrFC=0.1422]; Wilcoxon p<0.001, comparing noise with oscillations [mean ErrFC=0.2063] in **Figure 2H**). On the other hand, the fluctuating and oscillatory regimes better fit the functional connectivity than the noise regime. However, in this case, the fluctuating regime is significantly better than the oscillatory regime (Wilcoxon p<0.001, comparing fluctuations with oscillations box in **Figure 2H**).

We also evaluated the functional connectivity fitting for the same model-surrogates generated previously (see Methods) to compare with the corresponding optimal working point. The results clearly show significant differences with the obtained level of fitting with the optimal working point of the models (Wilcoxon p<0.001, comparing box fluctuations with *surr_fluct* and box oscillations with *surr_osc*).

Finally, in **Figure 2I**, we demonstrate the amplitude turbulence (the local Kuramoto parameter, R, see Methods) at the optimal fitting point of the three whole-brain model regimes contrasted with the empirical data (right subpanel) by rendering continuous snapshots for segments separated in time rendered on a flatmap of a brain hemisphere.

### Modelling results for less fine parcellation with 68 regions

Following the precise results of fitting the whole-brain model to the empirical data using a fine parcellation, we turned our attention to showing the fitting a less fine parcellation. We found that this was also not able to distinguish between fluctuating subcritical and oscillatory supercritical regimes. Specifically, we found that the level of fitting the empirical metastability defined as the standard deviation of the global Kuramoto order parameters is the same for both regimes.

We used during this second analysis a smaller brain parcellation, the Desikan-Killiany with 68 cortical regions of interest (ROIs), to be able to establish a node-level perturbative *in silico* protocol. We repeated the fitting procedure by exploring the parameter space (G,β) for the model in fluctuation supercritical and oscillatory subcritical regime. This parcellation is not suitable for computing amplitude turbulence, as is defined in Kawamura et al. (Kawamura et al., 2007) and Deco et al. (Deco & Kringelbach, 2020), due to the lack of spatial resolution. We thus fitted the metastability, which is the most similar measure computable in coarser parcellation (Deco, Kringelbach, et al., 2017). We found the pair (G,β) that minimizes the absolute difference between the empirical and simulated levels of metastability.

**Figure 3** shows the results of fitting the Hopf whole-brain model in the two different regimes (fluctuating and oscillatory, see upper row panel A and B) for the Desikan-Killiany parcellation (upper row panel C) in terms of functional connectivity and metastability. For each of these regimes, we defined a grid of the parameter space (G,β), and for each pair in the grid, we simulated 100 times the whole-brain dynamics, and we computed the level of fitting between the metastability (second row) and simulated and empirical FC (third row).

**Figure 3.**
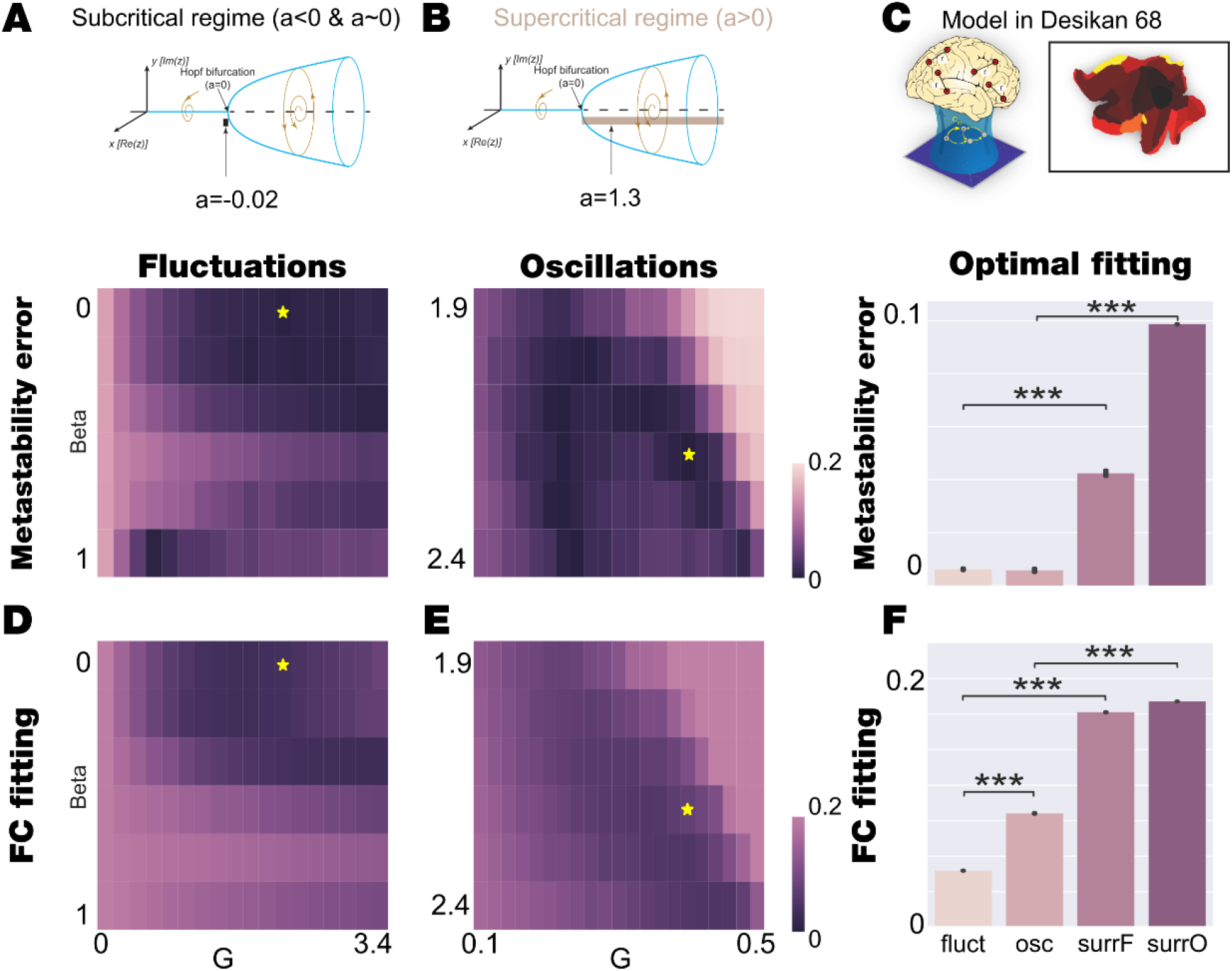
Model Desikan-Killiany fitting of fluctuations and oscillatory for 1) Metastability and 2) FC fitting in. **A-B)** We explored the bi-dimensional parameter space defined by β and G for fluctuating and oscillatory regime (bifurcation parameter a=-0.02 and a=1.3, respectively, indicated in the upper row) and computed the level of metastability error as the absolute difference between the empirical and simulated metastability. Yellow stars indicate the (β,G) combination that reaches the lowest metastability error in each regime. **C)** The upper subpanel shows the model fitting scheme procedure in coarser Desikan-Killiany parcellation. The bottom subpanel displays the barplot that indicates the statistical distribution of the metastability error obtained by simulating 20 trials with 100 subjects for each model regime with the parameters set at the corresponding working point. We also display the results of two model-based surrogates created by increasing the shear parameter of each model regime. The comparison between both model’s regimes at fitting the metastability shows that the two regimes are equally good (Wilcoxon, ns), while the rest of the comparisons are statistically significant (Wilcoxon, P<0.001). **D-E)** We explored the bi-dimensional parameter space defined by β and G for fluctuating and oscillatory regime computed the FC fitting as Euclidean distance between the empirical and simulated FC. Yellow stars indicate the (β,G) combination that reaches the lowest metastability error in each regime (the optimal working point obtained in panels A-B). **F)** The barplot indicates the statistical distribution of the FC fitting obtained by simulating 20 trials with 100 subjects for each model regime at the corresponding working point defined as the minimum metastability error. We also display the results of two model-based surrogates created by increasing the shear parameter of each model. All comparisons are statistically significant (Wilcoxon, P<0.001).

We found the optimal fitting for the level of metastability for each of the two regimes, indicated with a star in the second row of **Figures 3A-B. Figure 3A** shows the best fit for the fluctuating regime (a=- 0.02) with optimal (G,β)=(2.2,0) with minimal absolute difference between the simulated and empirical metastability M (here eM=4×10^−3^). **Figure 3B** shows the best fit for the oscillatory regime (a=1.3) with optimal (G,β)=(0.4, 2.2), which also produces an excellent fit with eD=1×10^−3^. We also computed the grid fitting for the FC, defined as the Euclidean distance between the simulated and empirical FC, for two regimes (**Figures 3D-E**), with a star in the grid indicating the optimal fit of metastability (which is the criterium for selecting the optimal working point since this is the most similar measure to turbulence). Note that these points do not correspond to the optimal fitting with FC in the two regimes.

The statistical comparison between the optimal working point of both model regimes (defined by the optimal fitting of the level of metastability, indicated with the stars in **Figures 3A-B**) is quantified in **Figure 3C and F**. We simulated 20 times the 100 repetitions of the whole-brain model at each regime working point and compare the level of metastability fitting and functional connectivity fitting (see Methods).

The results show that the fitting of the data by the fluctuating and oscillatory regimes is excellent (mean eM=6×10^−3^ in both cases) and not significantly different (Wilcoxon n.s, comparing first with the second bar in the second row of **Figure 3C**). On the other hand, the fluctuating regimes better fit the functional connectivity than the oscillatory regime (Wilcoxon p<0.001, comparing fluctuations with oscillations boxes in **Figure 3F**).

We also generated model-surrogates to compare with the corresponding optimal working point by setting the parameters of the model in the optimal working point but increasing β. Hence, we produced two surrogate models: *surr_fluct* for the fluctuating model surrogates using *a=-0*.*02* and (G,β) =(2.2, 3) and *surr_osc* for the oscillatory model surrogates using *a=1*.*3*. and (G,β) =(0.4, 3). The results clearly show significant differences comparing with the level of metastability fitting obtained by the optimal working point of the model in different regimes (Wilcoxon p<0.001, comparing fluctuation with *surr_fluct* box and oscillations with *surr_osc* boxes in the second row of **Figure 3C**). The same results were obtained for the fitting of the functional connectivity (Wilcoxon p<0.001, comparing fluctuation with *surr_fluct* box and oscillations with *surr_osc* boxes of **Figure 3F**).

The results also show that it could not distinguish between fluctuating subcritical and oscillatory supercritical regimes in coarser parcellation in terms of fitting the empirical data. We then focused our analysis on the perturbation response as an approach to disentangle between both models.

### Global strength-dependent perturbation distinguishes between fluctuating and oscillatory regimes

We implemented a global strength-dependent sustained perturbation that allows us distinguishing between fluctuating and oscillatory regimes for both the fine and coarse parcellations (**Figure 4**). We generated an *in silico* stimulus by adding an external periodic force applied equally to all nodes at the optimal working point in both model regimes (see Methods). We varied the strength of the external forcing *F*_*0*_ from 0 to 0.001 in steps of 0.0001, and for each amplitude, we simulated 100 times the perturbed and unperturbed model signals. We obtained the local and global Kuramoto order parameters (lKoP and gKoP) for the perturbed and unperturbed cases for the fine and coarse parcellation, respectively. We then computed the local and global Susceptibility and absolute Information Capacity as the mean and standard deviation of the subtraction between the perturbed and unperturbed lKoP and gKoP across trials (see Methods). We repeated this computation 20 times and **Figure 4C-F** shows the mean and standard deviation across repetitions. The subcritical fluctuating regime shows a rapid increase of the level of local Susceptibility in the fine parcellation (**Figure 4C**) and the level of global Susceptibility in the coarse parcellation (**Figure 4D**). The global absolute Information Capability also rapidly increase while the forcing strength increases (**Figure 4E-F** dark colors).

**Figure 4.**
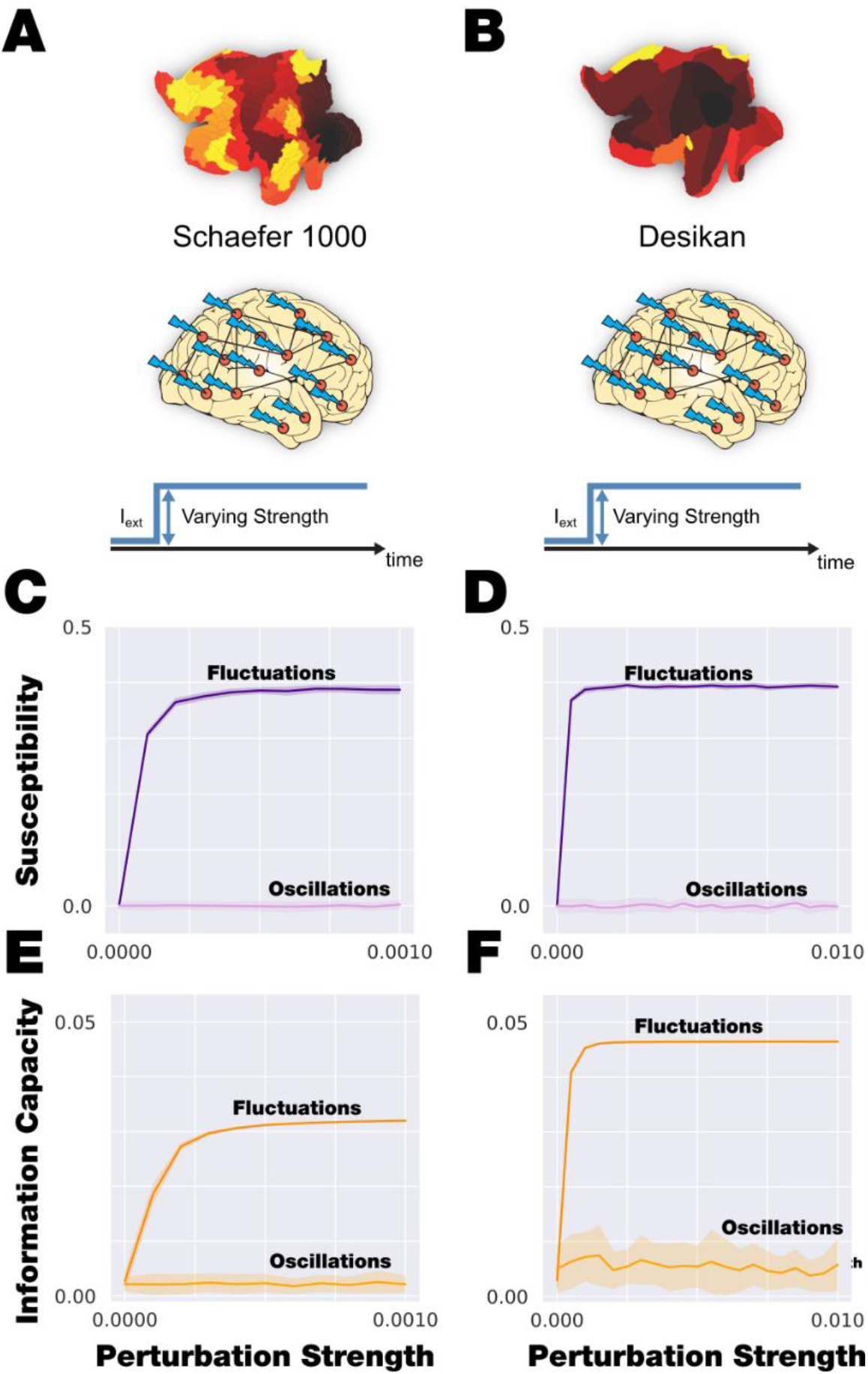
Global and sustained strength-dependent perturbation. **A)** We applied global strength-dependent, sustained perturbation in Schaefer1000 parcellation, and **B)** the same perturbation in Desikan-Killiany parcellation. **C-D)** The evolution of local and global Susceptibility (fine parcellation, panel C and coarse parcellation, panel D, respectively) as a function of perturbation strength. In dark purple is shown the response of the subcritical fluctuating regime, while in light purple, the behaviour of the supercritical oscillating regime. The subcritical regime is clearly more susceptible than the supercritical regime that is almost unaltered by the perturbation. **E-F)** The evolution of global absolute Information Capacity (fine parcellation, panel E and coarse parcellation, panel F, respectively) as a function of perturbation strength. In dark orange is shown the response of the subcritical fluctuating regime, while in light orange, the behaviour of the supercritical oscillating regime. The subcritical regime clearly changes the Information Capacity with the perturbation strength comparing with the supercritical regime that is almost unaltered by the perturbation.

In the supercritical oscillatory regime, the global Susceptibility and absolute Information Capability are constant in both parcellation along *F*_*0*_ (**Figure 4C-F** light colours). It is remarkable that the level of these measurements in this regime keep almost zero for all the strength forcing range, showing that the model in that regime do not respond under this global perturbation.

### Local strength-dependent perturbation also distinguishes between fluctuating and oscillatory regimes

We then used local strength-dependent sustained and non-sustained perturbations that also allows us distinguishing between fluctuating and oscillatory regimes (**Figure 5**). This is demonstrated using the less fine parcellation. This reduction of the number of regions allowed us to define a node-by-node perturbative approach. Firstly, we explored the model’s regime response by applying a sustained perturbation, and then we quantified the response to non-sustained external perturbation by the PCI.

**Figure 5.**
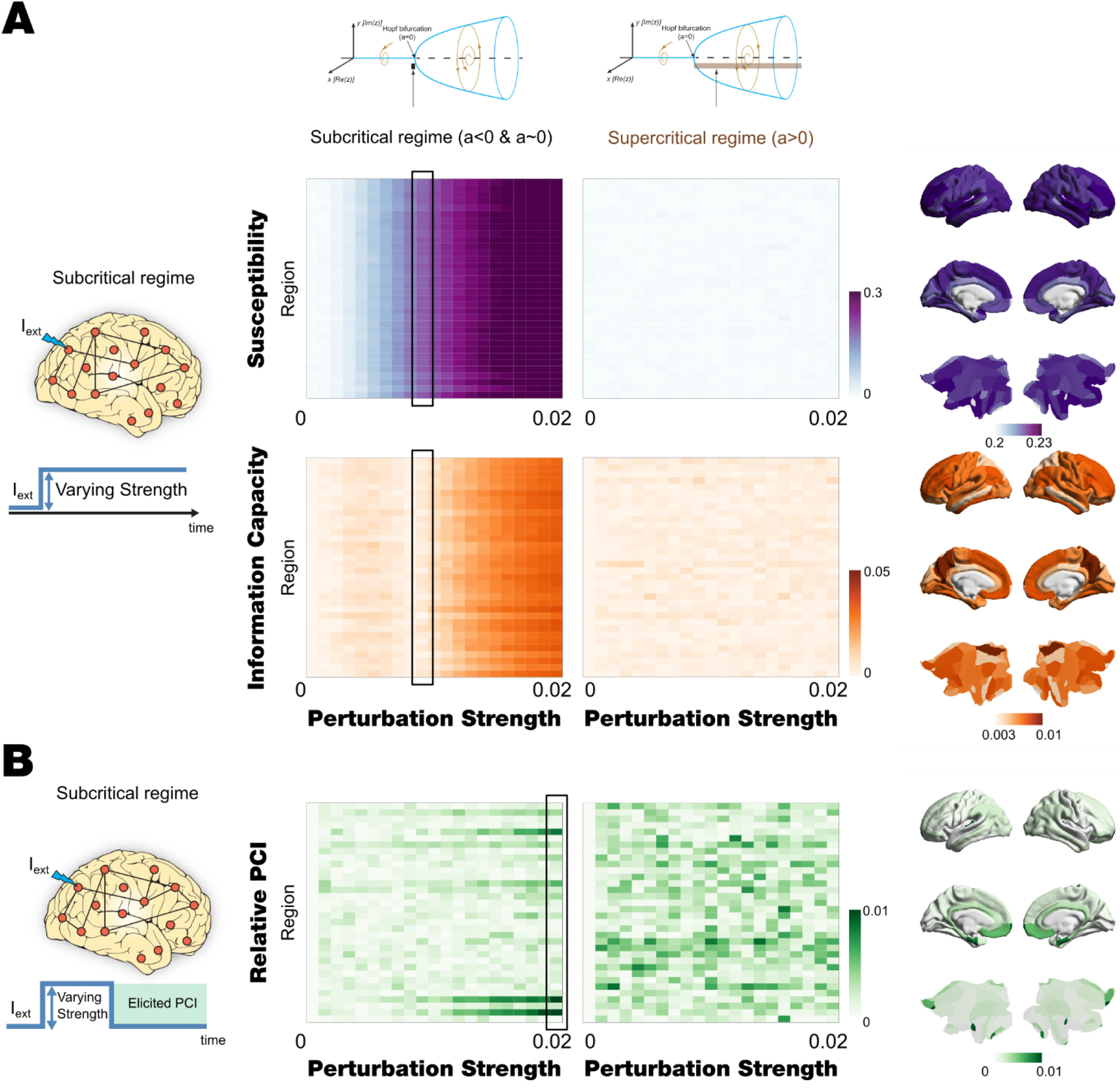
Local and Sustained/non-sustained strength-dependent perturbation. **A)** The evolution of Susceptibility (second row) and the absolute Information Capacity (third row) as a function of the perturbation strength and the perturbed pairs of homotopic nodes. The middle left panel displays the results for the subcritical regime (first row), and the middle right panel shows the response of the supercritical regime (first row). The right panels present the perturbative node hierarchy rendered onto the brain cortex for both measures (second and third row) for the case of a perturbation strength of 0.01 indicated with a box in middle left panel. **B)** Non-sustained PCI: The PCI is obtained by perturbing by pairs of homotopic nodes and different forcing amplitude. In the left column, the PCI results are obtained by perturbing the subcritical model in its corresponding working point with an external periodic force applied by pairs of homotopic nodes as a function of the amplitude of this forcing. In the right column, the same measurement is displayed but, in this case, for the supercritical model in its corresponding working point. The right panel shows the node-perturbative hierarchy in terms of PCI of each region for the maximum value of the forcing amplitude (indicated with black box in the middle-left panel) rendered onto a brain cortex.

### Susceptibility and Information Capability after local strength-dependent sustained perturbations

We systematically perturbed the model in each regime optimal working point by adding an external periodic force (see **Figure 5A** and Methods). We performed this *in silico* stimulation approach by forcing the 34 pairs of homotopic nodes (in the parcellation with 68 nodes) with forcing strength ranging from 0 to 0.02 in 0.001 steps. For each combination of nodes and amplitude, we ran 50 trials with 100 simulations, each computing the global Kuramoto Order parameter (gKoP) for the perturbed and unperturbed case (see Methods). We defined the node-level global Susceptibility and Information Capability as the mean and standard deviation across simulations of the subtraction between the node-perturbed and unperturbed gKoP, and we then averaged across trials. **Figure 5A** (second and third rows) shows the results for oscillatory (supercritical) and fluctuating (subcritical) regimes for both measurements. As in the global perturbation experiment, we noticed that the supercritical regime shows almost non-response under the perturbation, while the subcritical case presents variations across nodes and forcing amplitude. In this node-level perturbative approach, we can determine a hierarchy of perturbative effects by assessing node-by-node perturbation effect while the forcing amplitude increases. **Figure 5A** left panels shows a render onto brain cortex for both measurements in the subcritical regimen for 0.01 of forcing strength.

### PCI after local strength-dependent, non-sustained perturbations

We slightly modified our perturbative approach to bring the *in silico* stimulation protocol closer to *in vivo* experiments, as proposed by Massimini and colleagues (Casali et al., 2013). We simulated the external perturbation as an additive external force by pairs of homotopic nodes as in previous sections, but in this case, we focused on the response after the perturbation ends. Specifically, we simulated 600 volumes with the perturbation active, and we then evaluated the evolution of the signals in the following 200 volumes without perturbation. To investigate the behaviour of both model regimes, we adapted the PCI as is defined in Massimini et al. (Casali et al., 2013) to be applied on simulated BOLD signals (see Methods). This index gauges the amount of information contained in the integrated response to an external perturbation. **Figure 5B** displays the evolution of the PCI, computed as the normalised perturbed algorithmic complexity 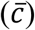 minus the background algorithmic complexity 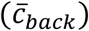, for both model regimes, for each pair of nodes and forcing strength. We found that in the oscillatory regime, the behaviour of the system after the perturbation is almost the same as the behaviour of the system without perturbation 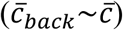, for all nodes and amplitudes (**Figure 5B** middle right panel). On the other hand, assessing the perturbation of the subcritical regime unveils a node hierarchy of the response under external perturbations (**Figure 5B** middle left panel).

These local responses under perturbations rendered onto the brain cortex for the maximal forcing amplitude is displayed in **Figure 5B** (right panel). It is remarkable that in the subcritical case, a set of nodes present the strongest response in terms of intensity (low values of PCI) and sensitivity (for lower forcing amplitudes). Most nodes present a moderate response for perturbations with forcing amplitude higher than 0.01, and other nodes remain unaltered.

### Regional heterogeneity and node-hierarchy perturbative organization

As shown in **Figure 6**, we also investigated how the node-hierarchy established in the previous section can be related to other sources of regional heterogeneity. We used different external sources of local heterogeneity, the T1w:T2w ratio and the principal component of transcriptional activity of an extensive set of specific brain genes (see Methods). We also compared with the anatomical and functional connectivity strength of each region, computed as 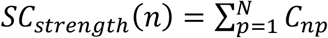 and 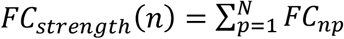 (well-known as Global brain connectivity, GBC), respectively, where C is the anatomical structural connectivity, and FC is the functional connectivity (see Methods). Finally, we compared the PCI node-hierarchy with the one found with global Susceptibility and Information Capability. We observed that the PCI hierarchical organisation is highly correlated with the other two perturbative measures obtained in the study. It is remarkable that an important difference between both experiments relies on that the first measures are computed after the perturbation while the second measures are computed after the perturbation.

**Figure 6:**
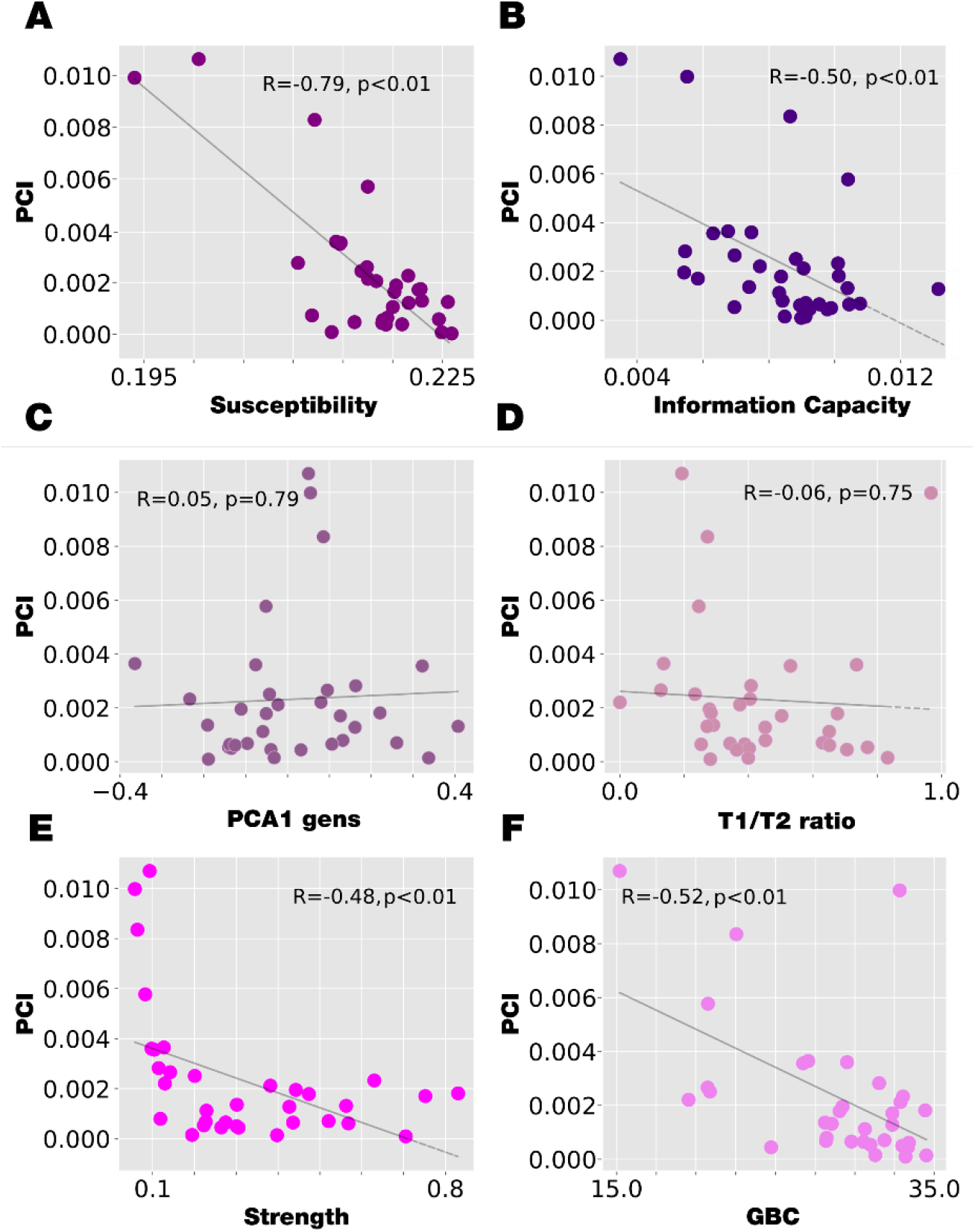
The correlation between the node-level PCI and other sources of regional-level heterogeneity. **A)** The correlation between node-level PCI and the node-level global Susceptibility is computed with significant negative correlation. **B) C)** The correlation between the node-level PCI and the first principal component of genes expression node information was computed with no correlation between variables. **D)** The same occurs in the correlation computed in blue circles between the node-level PCI and the ratio between the T1/T2 MRI. **E)** The correlation between the node-level PCI and the node functional connectivity strength (GBC) is computed obtaining a significant level of negative correlation. **F)** The correlation between the node-level PCI and the node anatomical strength is computed obtaining a significant level of negative correlation.

### Revealing the causal mechanistic principles of empirical results modulating resting state networks using stimulation

Experimental research has shown that TMS stimulation of specific brain regions can differentially modulate specific networks (Santarnecchi et al., 2018; Tik et al., 2017). We wanted to reveal the causal mechanistic principles and performed a network-level analysis testing the response of both model regimes (fluctuations and oscillations) using the Desikan-Killiany 68 parcellation, where each parcel belongs to one of the seven Yeo networks. The existing stimulation protocol was then applied in the same manner as in the previous analysis but now using a fixed forcing amplitude (*F*_0_ = 0.01). We computed the mean functional connectivity for the parcels belonging to each network using both fluctuating and oscillating regimes before and after the perturbation.

**Figure 7A** shows the seven Yeo resting state networks rendered on the medial and lateral surface of the right hemisphere of the brain. **Figure 7B** shows the differences between the perturbed and unperturbed FC for each model regime and the seven Yeo networks as a function of the perturbed node. We found that the fluctuating regime enhances the functionality for all perturbed nodes and all networks, while the oscillatory regimen is essentially unaltered. **Figure 7C** shows boxplots of the level of FC for each of the seven Yeo networks for the unperturbed and perturbed case for each model regime. We observed that not only the subcritical regime enhances the FC for all network but also allows representing different levels depending on the network. In contrast, the subcritical regime is almost constant for all networks.

**Figure 7.**
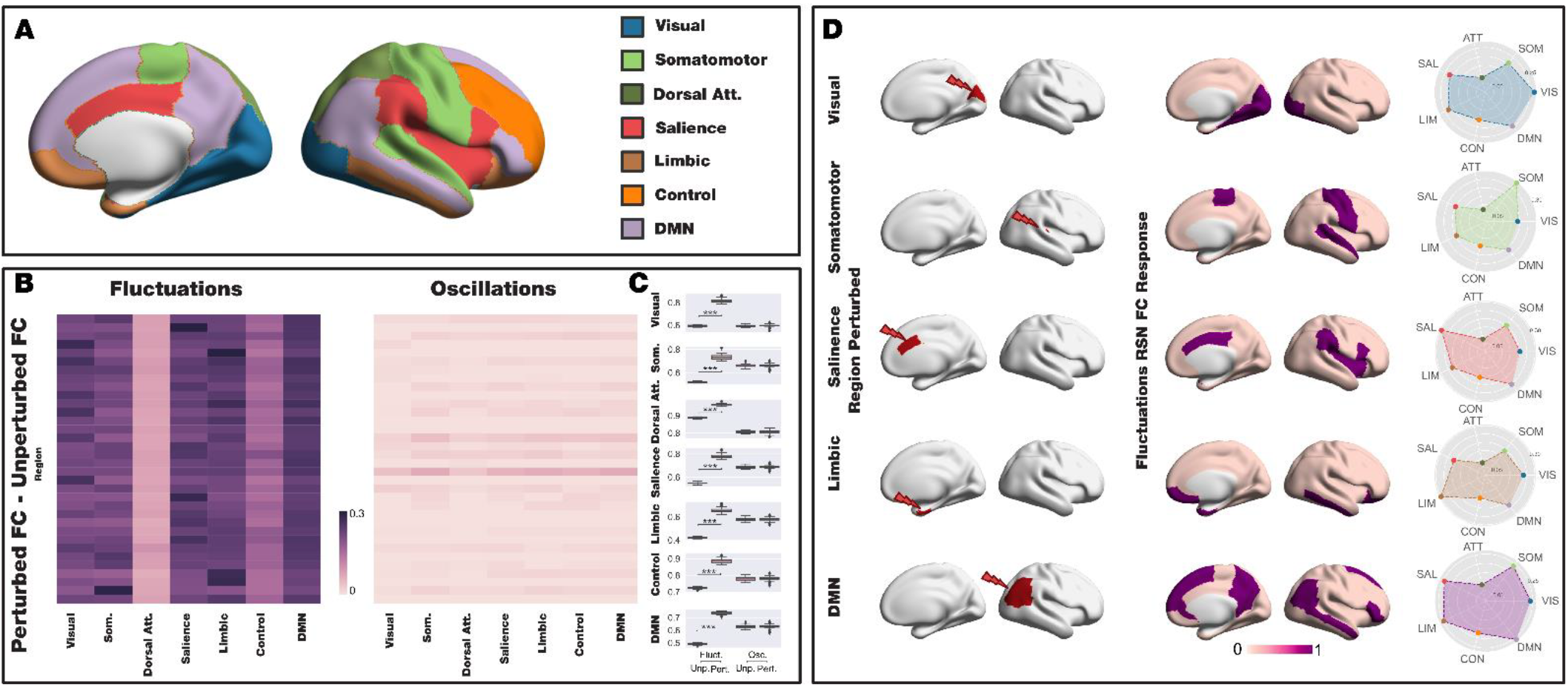
Revealing the causal mechanistic principles of empirical results modulating resting state networks using stimulation. **A)** For the reference, the seven Yeo resting state networks are rendered in the medial and lateral surface of the right hemisphere of the brain. **B)** Local and Sustained stimulation differentially enhances the resting state networks. The difference in the level of FC between the perturbed and unperturbed case is shown for the seven Yeo resting state networks as a function of the perturbed node. The subcritical regime enhances the FC for all networks and nodes, while the supercritical regime is much smaller and almost constant across nodes and networks. **C)** The seven subpanels show boxplots of the seven Yeo resting state networks with the FC of the two model regimes in the unperturbed and perturbed case. The subcritical regime shows higher levels for the 7 networks and while the supercritical case remains almost unaltered with the perturbation. The significance of the results was assessed using the Wilcoxon rank-sum test, where *** represents p<0.001. **D)** Left column shows a representative region (in red) in different resting state networks being perturbed in the fluctuating regime, which gives rise to a stabilisation of the respective network. The middle column is showing a rendering of the normalised difference between the perturbed and unperturbed activity, thresholded to top 15%. This can be seen in the spiderplots (right column), where the elicited activity is maximal for the stimulated network.

Given that we found that only the fluctuating regime is able to modulate the network after perturbation, we only used this regime to stimulate a representative region (in red) of different resting state networks (**Figure 7D**, left column). This gives rise to a stabilisation of the respective network. **Figure 7D** (middle column) shows renderings of the normalised difference between the perturbed and unperturbed activity, thresholded to top the 15% for five resting state networks, with the right column showing this change as spiderplots of the elicited activity for each seven Yeo resting state network. As can be seen, the stimulated network is also the most stabilised. This result demonstrates the underlying principles for the empirical findings of modulation of resting state network following stimulation. This provides crucial empirical evidence for the fluctuating regime.

## Discussion

Here we used whole-brain models to address a fundamental question in neuroscience of the origin of the fluctuations and oscillations found in global brain dynamics. We used strength-dependent perturbations of the whole-brain model fitting the empirical data to give a causal mechanistic description of human brain function, showing that this is generated by a delicate balance between fluctuation and oscillation on the edge of criticality. These results were obtained using the large-scale Human Connectome neuroimaging dataset of 1003 participants, which were subsequently used for massive computational whole-brain modelling studies. Crucially, the underlying causal mechanistic principles of empirically reported modulation of specific resting state network can be explained by stimulation in the fluctuating regime. Overall, the present strength-dependent perturbation framework demonstrates that maximal flexibility for the human brain comes from whole-brain dynamics poised on the edge between fluctuating and oscillatory regimes.

Specifically, we demonstrated that fluctuations and oscillations regimes of the Hopf whole-brain model are equally good at fitting the empirical data in terms of turbulence and functional connectivity representing asynchronous and synchronous background dynamics. We also demonstrated that strength-dependent *in silico* perturbations, either local or global, sustained or non-sustained provide valuable insights to unveil which regime is suitable for representing global brain dynamics and its capacity to encode external stimuli.

### Fluctuating and oscillatory regimes are distinguished by global strength-dependent perturbation

We found that global strength-dependent and sustained perturbation distinguishes between fluctuating and oscillatory regimes. The level of Susceptibility and Information Capability rapidly increase with amplitude strength in the subcritical fluctuating regime in fine-scale and coarser parcellations with 1000 and 68 regions, respectively. Conversely, the level of both measures in the supercritical oscillating regime remains almost constant along with the full range of amplitude strength. This result disentangled the two models, where the subcritical model clearly outperforms the supercritical model, providing a novel indication that the optimal dynamical behaviour is on the edge of criticality, between fluctuations and oscillations, as suggested by previous research (Deco et al., 2013; Deco, Kringelbach, et al., 2017; Spiegler et al., 2016).

Also considering the similarities with the thermodynamic phase transition and bifurcations in dynamical system (Bose & Ghosh, 2019), this result can be interpreted in the terms of the statistical criticality in brain dynamics. Previous research has demonstrated that the brain dynamics is poised near criticality, i.e., near the critical point of a phase transition (Chialvo, 2010; Cocchi et al., 2017), and at this point the system has the higher susceptibility, where a small perturbation is able to be propagated along the whole system. Following this comparison, we can claim that to be on the edge of the bifurcation is comparable to staying close to the critical point of the phase transition and the result of both scenarios is to amplify the effect of perturbation and thus increase complexity.

### Local strength-dependent perturbation can distinguish between dynamical regimes

Furthermore, it is also possible to investigate the model’s regime responses by applying local strength-dependent *in silico* perturbations. To this end, we used a less fine parcellation with 68 regions and systematically applied an external strength-dependent periodic force to all pairs of homotopic nodes. We also found that local strength-dependent and sustained perturbations efficiently discriminate between fluctuating and oscillatory regimes. We found that the node-by-node Susceptibility and Information Capacity increase with amplitude strength in the subcritical fluctuating regime, while in the supercritical regimen, both measures remain almost constant under the perturbations. Even more, in this node-level perturbative approach, we found a hierarchy of perturbative effect by assessing node-by-node response to the perturbation while the forcing amplitude increases in terms of Susceptibility and Information Capacity measures. These results extend the findings from previous research on elucidating the principles of deep brain stimulation (Saenger et al., 2017), transcranial direct current stimulation (Kunze et al., 2016) and recent research demonstrating in principle how to awaken a model of the sleeping brain (Deco et al., 2019; Perl et al., 2021) or how specific functional networks emerge after local stimulation (Spiegler et al., 2016, 2020).

### Local strength-dependent, non-sustained perturbations changes the PCI

Inspired by the pioneering results of perturbing the brain directly revealed by the empirical studies of Massimini and colleagues (Casali et al., 2013), we created a perturbative *in silico* strength-dependent local and non-sustained protocol which can provide testable empirical predictions in human participants by extending their use of PCI (Casali et al., 2013; Ferrarelli et al., 2010; Goldman et al., 2020; Rosanova et al., 2012). We found that in the oscillatory supercritical regime, the behaviour of the system after the perturbation is almost the same as without the perturbation for all nodes and amplitudes. The quantification of the response after the perturbation in the subcritical regime unveiled a node hierarchy of the response under external perturbations. As such, we were able to represent this hierarchy rendering onto the brain the value of the obtained PCI for each node at the maximal forcing amplitude.

### Hierarchical organisation revealed by perturbation of whole-brain model

We were able to reveal the hierarchical organisation through computing by PCI following local strength-dependent perturbations and comparing with other sources of regional heterogeneity. We used four of heterogeneity: 1) the myelination ratio (T1:T2w ratio), 2) the principal component of transcriptional activity of a large set of specific brain genes (Arnatkeviciute et al., 2019; Deco, Kringelbach, et al., 2021; Hawrylycz et al., 2012), 3) node-strength of structural and functional connectivity and 4) the hierarchies obtained for the Susceptibility and Information Capability computed for the local and sustained strength-dependent perturbations. We demonstrated that the PCI hierarchical organisation following local strength-dependent perturbations is highly correlated with the other two perturbative measures obtained in the study, which is correlated with the node-strength of structural and functional connectivity. Conversely, the PCI hierarchical organisation does not correlate with the T1:T2w ratio and PC1 of genes transcriptional activity.

These results show the power of perturbative *in silico* framework for addressing a fundamental question in neuroscience: namely, the role of the local fluctuations and oscillations in shaping the emergent global brain dynamical. By investigating the dynamics of the brain through a Hopf whole-brain model that allows switching from noisy asynchronous dynamics towards synchronous oscillations (Deco, Kringelbach, et al., 2017) we show that both dynamical regimes in microscopic and macroscopic scales are associated with different global brain states: while the first seems to be the dynamical background need to support a responsive brain state, the second is related to reduced states of consciousness (Goldman et al., 2019).

However, our framework is capable of distinguishing between both dynamical scenarios, but we also found that the perturbative hierarchy can provide an independent source of local information that can be used as *prior* in studies where heterogeneity plays a role (Deco, Kringelbach, et al., 2021; Kringelbach et al., 2020). These findings also pose a question regarding the relationship between each regional response capability to external stimuli and the role of fluctuations and oscillations. Future research could investigate heterogeneous models that allow each region to be in fluctuating or oscillatory regimes, the causal link between the local model regime and whole-brain susceptibility. Ultimately, this could help cast new light on the mechanistic interpretation of the local dynamics responsiveness in terms of the global response (Destexhe & Contreras, 2006).

### Empirical evidence for the fluctuating regime

Finally, we evaluated the ability of stimulation in the fluctuating regime for modulating resting state networks by studying the functional connectivity over the entire network. This was inspired by experimental results demonstrating that different stimuli can bring about network-specific modulation (M. D. Fox et al., 2012; Santarnecchi et al., 2018; Tik et al., 2017). The local and sustained stimulations in the whole-brain model may approximate invasive stimulation techniques such as deep brain stimulation, DBS (Kringelbach et al., 2007), as well as non-invasive stimulation techniques such as transcranial magnetic stimulation, TMS (Dayan et al., 2013). We compared the dynamically responsive networks to external stimulus in different brain targets for both model regimes. In fluctuating/subcritical regime, the brain reacts to specific local stimulation with activity patterns that closely mimic the seven Yeo resting state networks (Yeo et al., 2011). We found that perturbing a region in a given resting state network led to stabilisation of that network. Our results are aligned with previous computational studies demonstrating that perturbative approaches are able to predict empirical observation, such as the emergence of large-scale functional networks (Spiegler et al., 2016; Yang et al., 2021). Importantly, and supporting the superiority of the subcritical, fluctuating regime, in the supercritical oscillation regime the brain response remains almost unaltered when perturbing all nodes in the seven Yeo networks. In summary, preserving the resting state network structure is better represented by the subcritical regime, showing that dynamically responding brain networks are the outcome of a model poised on the edge of bifurcation.

### Challenges and opportunities for in silico perturbation approaches

The findings have been made possible by the whole-brain modelling framework developed over the last decade (Breakspear, 2017; Deco, Kringelbach, et al., 2017; Honey et al., 2009; Jirsa et al., 2002) A clear advantage of using such data-constrained whole-brain models is its potential use for studying stimulation protocols, as this enables an exhaustive search and optimization of all underlying parameters and locations *in silico*, and it may offer insights into the self-organization of widespread networks (Deco et al., 2018, 2019). This strategy allowed to computational assess the stimulation-induced transition between brain states as an insight of treatments prognosis (Kringelbach & Deco, 2020), awakening from sleep stages (Deco et al., 2019), or defined perturbative metrics as a brain state characterization (Perl et al., 2021).

Nevertheless, despite there is much empirical evidence that clearly reflects the change of dynamics following perturbations (Angelakis et al., 2014; Ozdemir et al., 2020; Zhang & Song, 2017), and the computational *in silico* results are really promising, the field awaits to confirm the whole-brain modelling predictive power. A potential path to doing such experiments could come from generative whole-brain models of the brain activity in animals (including non-human primates) (Deco et al., 2014; Ponce-Alvarez et al., 2021; Shen et al., 2019) that allow performing both models and empirical tests (Yang et al., 2021). In the future, these models could be used for investigating the changes in brain state between awake and anaesthesia non-human primates (Barttfeld et al., 2015), and suggest potential stimulation sites for transitioning between brain states, which can then be directly probed in these animal models. In this work, we pursued a novel approach by joining experimental and computational approaches. Our findings point to the possibility of strategically defined synthetic brain stimulations close to the specific experiments as an extension of the PCI (Casali et al., 2013; Goldman et al., 2020).

Overall, here we have shed further light on a long-standing question in neuroscience, namely how and why brain states are characterised by complex, fluctuating and oscillating dynamics. Our results provide new evidence using strength-dependent perturbations of the whole-brain model, revealing that brain function emerge at a delicate balance between fluctuation and oscillation on the edge of criticality.

## Methods

### Neuroimaging ethics

The Washington University–University of Minnesota (WU-Minn HCP) Consortium obtained full informed consent from all participants, and research procedures and ethical guidelines were followed in accordance with Washington University institutional review board approval.

### Neuroimaging Participants

The data set was obtained from the Human Connectome Project (HCP) where we chose a sample of 1000 participants during resting state. The full informed consent from all participants was obtained by The Washington University–University of Minnesota (WU-Minn HCP) Consortium and research procedures and ethical guidelines were followed per Washington University institutional review board approval.

### Brain parcellations

To compute the empirical and simulated level of turbulence in brain dynamics defined as Deco et al. (Deco & Kringelbach, 2020), we used the publicly available population atlas of cerebral cortical parcellation created by Schaefer and colleagues (Schaefer et al., 2018). They provide several parcellations sizes available in surface spaces, as well as MNI152 volumetric space. We used the Schaefer parcellation with 1000 brain areas, estimated the Euclidean distances from the MNI space, and extracted the timeseries from the HCP surface space version.

Desikan and colleagues created an automated labelling system subdividing the human cerebral cortex into standard gyral-based neuroanatomical regions identifying 34 cortical ROIs in each hemisphere (Desikan et al., 2006). In the second section of this work, we used this parcellation to assess systematically the perturbation protocol ROI by ROI.

### Neuroimaging acquisition for fMRI HCP

The HCP web (http://www.humanconnectome.org/) provides the complete details for the acquisition protocol, participants information, and resting-state data. We used one resting-state acquisition of approximately 15 minutes, acquired for 1003 HCP participants scanned on a 3-T connectome-Skyra scanner (Siemens).

### Preprocessing and extraction of functional timeseries in fMRI resting data

The resting-state data were preprocessed using FSL (FMRIB Software Library), FreeSurfer, and the Connectome Workbench software (Smith et al., 2013) as reported in (Deco, Perl, et al., 2021), which is described in detail on the HCP website. Briefly, the preprocessing included correction for head motion, spatial and gradient distortions, intensity normalisation and bias field removal, registration to the T1-weighted image, transformation to the 2mm Montreal Neurological Institute (MNI) space, and FIX artefact removal (Schröder et al., 2015; Smith et al., 2013). Artefactual components were removed by using ICA+FIX processing (Independent Component Analysis followed by FMRIB’s ICA-based X-noiseifier (Griffanti et al., 2014)). Preprocessed timeseries of all grayordinates are in HCP CIFTI grayordinates standard space, and available in the surface-based CIFTI file for each participant.

Custom-made Matlab scripts were applied using the ft_read_cifti function (Fieldtrip toolbox (Oostenveld et al., 2011)) to extract the timeseries of the grayordinates in each node of the Schaefer parcellation. Furthermore, the BOLD timeseries were transformed to phase space by filtering the signals within the range 0.008-0.08 Hz (M. Fox, 2005), and the low-pass cut-off to filter the physiological noise, which tends to dominate higher frequencies (Cordes et al., 2001; M. Fox, 2005).

### Structural connectivity using dMRI

The structural connectivity was obtained from the Human Connectome Project (HCP) database, which contains diffusion spectrum and T2 weighted imaging data from 32 participants. The acquisition parameters are described in detail on the HCP website. Briefly, the neuroimaging data were processed using a generalised q-sampling imaging algorithm developed in DSI studio (http://dsi-studio.labsolver.org). A white-matter mask was estimated by segmenting the T2-weighted images and images were co-registered to the b0 of the diffusion data by using SPM12. In each participant, 200,000 fibres were sampled within the white-matter mask. Fibres were transformed into MNI space using Lead-DBS (Horn & Blankenburg, 2016). We used the standardised methods in Lead-DBS to produce the structural connectomes for both Schaefer 1000 parcellation (Schaefer et al., 2018) and Desikan-Killiany 68 parcellation (Desikan et al., 2006), where the connectivity was normalised to a maximum of 0.2. The preprocessing implemented is freely available in the Lead-DBS software package (http://www.lead-dbs.org/) and is described in detail by Horn and colleagues (Horn et al., 2017).

### Whole-Brain Model

Whole-brain models have been used during the last decade to describe the most important features of brain activity. These models provide an optimum balance between complexity and realism, based on the fact that despite the macroscopic collective brain behaviour is an emergent of millions of smalls units interacting endowed with independent properties. One of the macroscopic dynamical features is that the collective behaviour dynamics can range from fully synchronous to stable asynchronous state governed by random fluctuations. The simplest dynamical system capable of presenting both behaviours is the one described by a Stuart Landau non-linear oscillator, which is mathematically described by the normal form of a supercritical Hopf bifurcation:

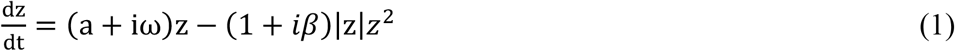

Where *z* is a complex-valued variable (*z* = *x* + *iy*), ω is the intrinsic frequency of the oscillator, and β is the shear factor. The bifurcation parameter *a* changes qualitatively the nature of the solutions of the system, if *a>0* the system engage in a limit cycle and presents self-sustained oscillations so-called the supercritical regime and when *a<0* the dynamics decay to a stable fixed point so-called the subcritical regime (Fig. 1E).

The coordinated dynamics of the resting state activity are modelled by introducing coupling between these oscillators. Previous research has demonstrated that whole-brain models based on Stuart Landau oscillators ruling the local dynamical behaviour have the capability to describe the time average behaviour (static functional connectivity) and dynamical behaviour (functional connectivity dynamics – FCD) on brain dynamics when the coupling between the oscillators is determined by the structural connectivity (Deco, Kringelbach, et al., 2017; Hansen et al., 2015; Jobst et al., 2017). Here, based on recent work, we assume that the coupling is determined by a combination of the *exponential distance rule* (EDR) and the long-range connection present in the structural connectivity (EDR-LR) (Deco, Perl, et al., 2021). The mathematical expression that rules this coupling factor is:

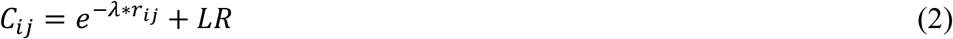

Where, λ stands for the exponential space decay fitted from empirical data and fixed at λ=0.18 mm^-1^ (Deco & Kringelbach, 2020), *r*_*ij*_ is the Euclidian distance between the node *i* and *j* and LR are the long-range connections extracted from the anatomical structural connectivity. The dynamical of the region (node) *i* in the coupled whole-brain system is described in cartesian coordinates:

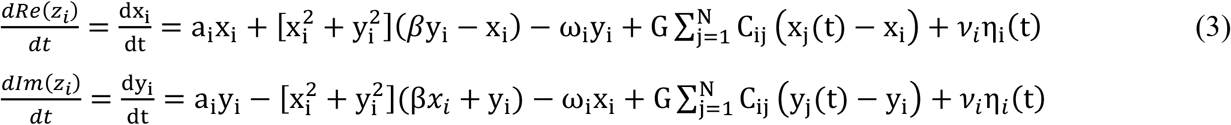

Where *η*_*i*_*(t)* is an additive Gaussian noise with standard deviation *ν* and G is a factor that scales the coupling strength equally for all the nodes. This whole-brain model has been shown to reproduce essential features of brain dynamics observed in different neuroimaging recordings (Deco, Kringelbach, et al., 2017; Piccinini et al., 2021) in the subcritical regime (i.e., *a<0*) and no shearing effect (β=0).

### Model optimal working point in (β,G) parameter space and regime comparison

We incorporate the shear factor as a global fitting parameter and the global scaling factor (G). In the first part of this study, we fit the level of turbulence using the Schaefer 1000 parcellation. We perform an exhaustive exploration of the parameter space (β,G), seeking the optimal working point of the model in noise regime (a=-1.3), subcritical regime (a=-0.02) and supercritical regime (a=1.3). In the supercritical case, we explore a grid of β=[1.7; 2.5] and G=[0.13; 0.22] in 0.1 steps, whereas in the noise and subcritical case we explore a grid of β=[0; 0.8] and G=[0.45; 1.8] in steps of 0.1 and 0.15, respectively. We generate 100 simulations with the same number of volumes (1200 volumes) and sampling rate (0.72 s) of empirical data for each pair (β,G) on the grid and compute the simulated level of turbulence and functional connectivity as the Pearson correlation between nodes signals. We estimated the fitting of level of turbulence as the absolute value of the difference between the average of the empirical and simulated level of turbulence and the functional connectivity fitting as the Euclidean distance between the empirical and simulated FC.

For comparing how good each regime is at fitting the empirical data, we generate 100 simulations with the same number of volumes (1200 volumes) and sampling rate (0.72 s) as the empirical data at the optimal working point of the three regimes. We compute the error of turbulence fitting and the FC fitting as the average value across simulations. We repeat 20 times each set of simulations. We also reproduce the same amount of simulation for two model surrogates consisting in increase to 6 the value of the shear parameter (β) for the optimal working point of the subcritical and supercritical regime.

In the second part of this study, we change the parcellation to the Desikan-Killiany parcellation with 68 nodes. The advantage of this parcellation is that it allows us to establish a systematical perturbation protocol at node scale within affordable computational time. The disadvantage is that the level of turbulence definition, as in Deco et al. (Deco & Kringelbach, 2020), is not computable in small parcellations, i.e., low distance resolution. We fit the level of metastability by computing the absolute difference between the empirical and simulated brain signals. We perform an exhaustive exploration of the parameter space (β,G), seeking the optimal working point of the model only in supercritical regime (a=1.3) and subcritical regime (a=-0.02), we discarded the noise regime in light of the results obtained in the fine parcellation analysis. In the supercritical case, we explore a grid of β=[1.9; 2.4] and G=[0.1; 0.5] in 0.1 and 0.02 steps respectively, while in the subcritical case we explore a grid of β=[0; 1] and G=[0; 3.4] in steps of 0.2 both. We generate 100 simulations with the same number of volumes (1200 volumes) and sampling rate (0.72 s) of empirical data for each pair (β,G) on the grid and compute the simulated level of metastability and the functional connectivity fitting using the Euclidean distances between the empirical and simulated FC (see Methods below). For comparing how good each regime is at fitting the empirical data, we generate 100 simulations with the same number of volumes (1200 volumes) and sampling rate (0.72 s) as the empirical data at the optimal working point of the three regimes. We compute the error of metastability fitting and the FC fitting as the average value across simulations. We repeat 20 times each set of simulations. We also reproduce the same amount of simulation for two model surrogates, which increase to 3 the value of the shear parameter (β) for the optimal working point of the subcritical and supercritical regimes.

### Perturbative in silico protocol

We model an external oscillatory perturbation and investigate the response of the whole-brain model fitted to the aforementioned observables in each parcellation in different model regimes. The stimulus was represented as an external additive periodic forcing term, given by *F*_*j*_ = *F*_0*j*_ cos(*ω*_*j*_*t*) + *iF*_0*j*_ sin(*ω*_*j*_*t*), in the corresponding real and imaginary part of the node *j* equation (eq. 3). The purpose of this perturbation was to model the effects of external stimulation (TMS, tACS). In the first part of this study, we simulate a global strength-dependent, sustained perturbation by applying the external forcing equally for all nodes (F_0_) at the node’s empirical frequency average (ω). We vary the forcing strength (F_0_) from 0 to 0.001 in 0.0001 steps. We then generate 50 trials with 50 simulations each one for each step and compute the perturbed and unperturbed local (global) Kuramoto order parameter in the fine (coarse) parcellation. Finally, we assess the behaviour of each model regime using the computation of local Susceptibility and Information capability in fine parcellation and through the global Susceptibility and local Information capability in coarser parcellation (see Methods below).

In the second part of the study, we simulate local strength-dependent, sustained and non-sustained perturbations adding an external periodic force by pairs of homotopic nodes. In this way, we obtain 34 *in silico* experiments varying the amplitude of the force, *F*_*0*_, from 0 to 0.02 in steps of 0.005 and generating 50 trials with 100 simulations each one for each step.

We assess the model’s response to the sustained perturbation by computing the global Susceptibility and Information Capacity by pairs of homotopic nodes and amplitude. We assess the model’s response to the non-sustained perturbation through computing the perturbative complexity index (PCI) for each forcing amplitude and pair of perturbed nodes.

### Measure of amplitude turbulence

The level of amplitude turbulence measure comes from the seminal studies by Kuramoto investigating turbulence in coupled oscillators (Kuramoto, 1984) and by Deco and Kringelbach that applied this concept to whole-brain dynamics (Deco & Kringelbach, 2020). Specifically, in a coupled oscillator framework, the Kuramoto local order parameter (lKoP) represents a spatial average of the complex phase factor of weighted coupling of local oscillators. The modulus of the Kuramoto local order parameter (*R*_*n*_(*t*)) is considered a measure of the local level of synchronization and is computed as:

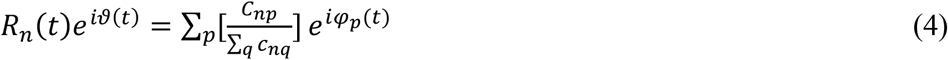

where φ_p_ is the phase of the BOLD signal of the node *p* and *C*_*np*_ is the strength of the coupling between node *n* and *p* determined by the exponential distance rule (first term of Eq. 2). We then compute the amplitude turbulence, D, as the standard deviation across time and space of the modulus of the lKoP, *R*_*n*_*(t)*:

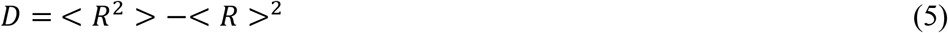

where brackets stand for the average across time and space.

We computed the error in fitting the level of turbulence (eD) as the absolute value of the difference between the empirical and simulated level of turbulence:

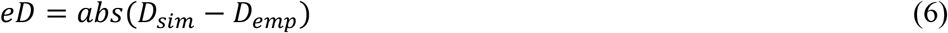

### Measure of metastability

The level of metastability measure was implemented in previous research to characterize the dynamics of the fluctuations in brain activity in different brain states (Deco, Kringelbach, et al., 2017; Jobst et al., 2017; Piccinini et al., 2021). Briefly, the metastability denotes the variability of the global synchronization as measured by the Kuramoto order parameter (gKoP), *gR(t)*,:

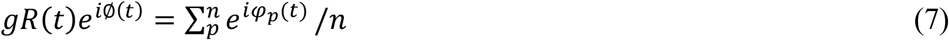

where φ_p_ is the phase of the BOLD signal of the node *p* and *n* is the total number of nodes in the parcellation. Thus, the metastability is the standard deviation of *gR(t)* across time:

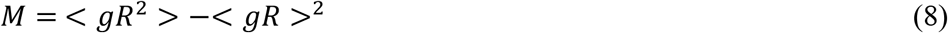

The fitting of metastability is defined as the absolute difference between the empirical and simulated level of metastability:

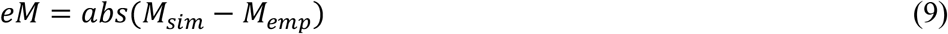

### Measure of Susceptibility

We define the whole-brain model susceptibility as the brain’s sensitivity to the processing of external periodic stimulations. We perturb the Hopf model in the supercritical and subcritical regime by adding an external periodic force with different amplitudes (see Methods, perturbative in silico protocols). We estimate the sensitivity of the perturbations on the spatiotemporal dynamics following previous work, which determines the susceptibility in a system of coupled oscillators based on the response of the Kuramoto order parameter (Daido, 2015). In the first part of this study, we extend this concept by assessing the variability of the modulus of the local Kuramoto order parameter, i.e., 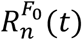 for the perturbed case for each value of forcing amplitude (*F*_*0*_), and *R*_*n*_(*t*) for the unperturbed case. We define local susceptibility in the following way:

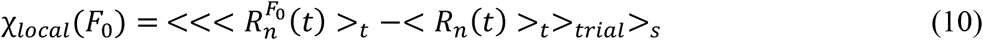

where <>_t_, <>_trials_ and <>_s_ are the mean averages across time, trials, and space, respectively.

In the second part of the study, we estimate the sensitivity of these perturbations by measuring the modulus of the global Kuramoto order parameter (gKoP), *gR(t)*, as a measurement of the global level of synchronization of the *n* nodes signal (Deco, Kringelbach, et al., 2017):

We compute the gKoP 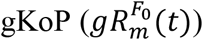 for the perturbed case for each value of forcing amplitude (*F*_0_) and pairs of perturbed nodes (*m*) and *gR*(*t*) for the unperturbed case. We define the global Kuramoto order parameter and global Susceptibility as follows:

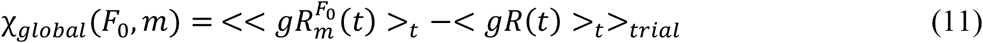

where <>_t_, <>_trials_ are the mean averages across time and trials.

### Measure of Information Capability

We define the Information Capability as a measure to capture how different external stimulations are encoded in the dynamics. We perturb the model in both regimes as above and compute for the first part of the study the perturbed and non-perturbed local Kuramoto order parameter for each forcing amplitude and, for the second part, the global Kuramoto order parameter for each forcing amplitude and perturbed nodes. The analytical computation of the Information Capability is through the standard deviation across trials of the difference between the perturbed Kuramoto order parameters and unperturbed ones. For the first part of the study, when we compute the local Kuramoto order parameter is computed as follows:

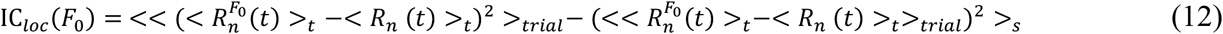

For the second part of the work, we compute the global Kuramoto order parameter and perturb by pairs of homotopic nodes (*m*) at different forcing amplitude (*F*_*0*_):

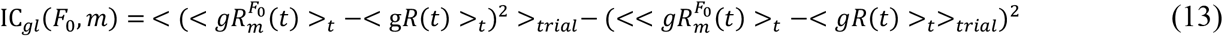

We then define absolute Information Capability (aIC) as the absolute difference between the IC at each forcing strength and the IC at zero-forcing for the global and local.

### Measure of PCI

We compute the perturbation complexity index (PCI) following the study of Casali and colleagues (Casali et al., 2013), where they implemented this index to characterise the empirical response to external stimuli in different states of consciousness. We simulate the perturbation described above by an external periodic force applied by pairs of homotopic nodes with different forcing amplitudes. For each case, we generate 100 simulations with 800 volumes and a sampling rate (0.72 s) for the optimal working point of each model regime. The first 600 volumes with the external force perturbing the system and the last 200 volumes without the perturbation. :

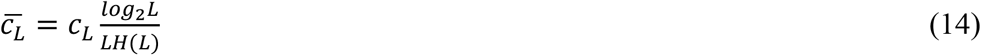

where *c*_*L*_ is the Lempev-Ziv complexity as a measure of algorithmic complexity (Lempel & Ziv, 1976), *L* is the length of the binary sequence, and *H(L)* is the source entropy of a sequence of length *L* that normalise the measure in order to be 1 to random sequences. For this purpose, we create a binary spatiotemporal distribution by z-scored the simulated times series after perturbation, *ts(n,t)*, where if *ts*_*zscore*_*(n,t)>2=1* and if *ts*_*zscore*_*(n,t)<2=0*. We then average across simulations the computed PCI for each pair of nodes and each forcing amplitude. To assess the response under external perturbation of both regimens, we compared the computed value after perturbation, 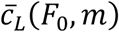, with the background level computed over simulated signal in each regime working point without perturbation 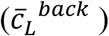

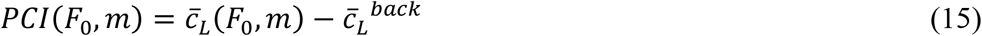

### Regional heterogeneity data

Here, we use different sources of regional heterogeneity to compare with the node-level hierarchy establish by the perturbation response. We consider the ratio T1w:T2w, which is sensitive to myelin content (Burt et al., 2018) and the first principal component (PC1) of transcriptional activity for 1,926 brain-specific genes. To this end, we use data from the Allen Institute Human Brain Atlas (AHBA), which comprises microarray data quantifying the transcriptional activity of >20,000 genes in >4,000 different tissue samples distributed throughout the brain, taken from six post-mortem samples. The AHBA data were processed following the pipeline developed in Arnatkevicuite et al. (Arnatkeviciute et al., 2019). To adapt the gene expression information into node-level heterogeneity information to Desikan-Killiany parcellation, we used the same approach explained in previous work (Deco, Kringelbach, et al., 2021).

### Statistical Analyses

Differences in model fits to empirical properties, as well as the resting state network enhancement, were assessed using pairwise Wilcoxon rank sum tests. The significance of each model regime fitting was assessed by comparing with model surrogates.

## Acknowledgments

YSP, AE and GD are supported by HBP SGA3 Human Brain Project Specific Grant Agreement 3 (grant agreement no. 945539), funded by the EU H2020 FET Flagship programme.

## Author Contributions

Y.S.P. designed the study, developed the methods, performed analyses, and wrote and edited the paper. M.L.K and G.D. designed and supervise the study, developed the methods, wrote and edited the paper. A.E. and E.T. helped with methodology revision, results discussion and edited the paper.

